# Optogenetic tools for public goods control in *Saccharomyces cerevisiae*

**DOI:** 10.1101/2021.06.28.450270

**Authors:** Neydis Moreno Morales, Michael T. Patel, Cameron J. Stewart, Kieran Sweeney, Megan N. McClean

## Abstract

Microorganisms live in dense and diverse communities, with interactions between cells guiding community development and phenotype. The ability to perturb specific intercellular interactions in space and time provides a powerful route to determining the critical interactions and design rules for microbial communities. Approaches using optogenetic tools to modulate these interactions offer promise, as light can be exquisitely controlled in space and time. We report new plasmids for rapid integration of an optogenetic system into *Saccharomyces cerevisiae* to engineer light-control of expression of a gene of interest. In a proof-of-principle study, we demonstrate the ability to control a model cooperative interaction, namely the expression of the enzyme invertase (SUC2) which allows *S. cerevisiae* to hydrolyze sucrose and utilize it as a carbon source. We demonstrate that the strength of this cooperative interaction can be tuned in space and time by modulating light intensity and through spatial control of illumination. Spatial control of light allows cooperators and cheaters to be spatially segregated, and we show that the interplay between cooperative and inhibitory interactions in space can lead to pattern formation. Our strategy can be applied to achieve spatiotemporal control of expression of a gene of interest in *Saccharomyces cerevisiae* to perturb both intercellular and interspecies interactions.

**Importance:** Recent advances in microbial ecology have highlighted the importance of intercellular interactions in controlling the development, composition and resilience of microbial communities. In order to better understand the role of these interactions in governing community development it is critical to be able to alter them in a controlled manner. Optogenetically-controlled interactions offer advantages over static perturbations or chemically-controlled interactions as light can be manipulated in space and time and doesn’t require the addition of nutrients or antibiotics. Here we report a system for rapidly achieving light-control of a gene of interest in the important model organism *Saccharomyces cerevisiae* and demonstrate that by controlling expression of the enzyme invertase we can control cooperative interactions. This approach will be useful for understanding intercellular and interspecies interactions in natural and synthetic microbial consortia containing *Saccharomyces cerevisiae* and serves as a proof-of-principle for implementing this approach in other consortia.

## Introduction

Interactions between individual cells and species dictate the development and phenotype of microbial communities^1–3^. These interactions are regulated in time and space, and often arise due to the different metabolic capabilities of specific cells and species^4,5^. Cooperative interactions are common, and cooperativity is often characterized by the presence of a shared public good which is produced by cooperative cells (producers) and freely available to other cells^3,6,7^. Production of the public good is often costly, and cooperative interactions are susceptible to the presence of “cheaters”, cells which exploit the public good without providing any contribution of their own^8^.

The budding yeast *Saccharomyces cerevisiae* engages in a cooperative interaction by secreting invertase, an enzyme which catalyzes the hydrolysis of sucrose into glucose and fructose. Due to its long domestication history and early enzymatic research on invertase^9–11^, invertase secretion by *S. cerevisiae* has long been used as a model system for studying public goods interactions and the emergence of cooperation in microbial communities. The *S. cerevisiae* genome contains several unlinked loci encoding invertase (SUC1-SUC8)^12,13^ but all except SUC2 are located within telomere sequences^14^. The strain used in this study (S288C) encodes only one functional invertase enzyme, SUC2^12,15^. There is a constitutively expressed intracellular form of invertase, but the secreted, glycosylated form which is regulated by glucose repression and important for cooperativity is secreted into the periplasmic space^16^. Most invertase (95%) remains in the cell wall; nevertheless yeast capture only a small fraction of the sugars that sucrose hydrolysis releases with most of the glucose and fructose diffusing away to be utilized by other cells^17–19^.

Hence the sugars produced from sucrose hydrolysis represent a “public good”. Invertase is costly to produce, and producing populations are susceptible to invasion by cheaters^17,20^.

There is growing evidence from both experiments and simulations that when and where a public good is produced within a microbial community can have dramatic consequences for community stability and the maintenance of cooperativity^21–27^. The spatial arrangement of genotypes within microbial communities can influence whether or not producers sufficiently benefit from the production of public goods, or whether cheaters are able to invade and take-over the community^3,26,28–30^. Indeed, efficient use of public goods has been identified as a possible driver for the evolution of multicellularity^31^. Furthermore, the dynamic control of public goods in both space and time could be used to manipulate synthetic consortia for applications in bioproduction and biotechnology^32,33^. Yet few tools exist for spatiotemporal control of specific community interactions.

Optogenetic tools offer the potential to overcome this limitation by utilizing genetically encoded light-sensitive proteins to actuate processes within the cell in a light-dependent manner. Light is a powerful actuator as it is inexpensive, easily controlled in time and space, and *S. cerevisiae* contains no known native photoreceptors^34^. Light can be rapidly added and removed from cell cultures or spatially targeted^35– 38^, meaning it can be used to study how regulation of microbial interactions determines microbial community development^39–41^. We report here the development of an optogenetic tool that allows the expression of a specific metabolic enzyme of interest to be put under light control in *S. cerevisiae*. Using this system, we demonstrate that we can use light to control when and where invertase is expressed within well-mixed and spatially organized populations of *S. cerevisiae*. Light control of this cooperative interaction shows that invertase expression in a community of yeast has important effects on overall community growth and spatial structure. Our results suggest that optogenetic control of microbial interactions is an important new approach to understanding and engineering microbial communities.

## Results

### Plasmid design and system overview

To enable light-based control of cooperativity we first developed constructs that, when integrated into yeast, allow us to make expression of a specific gene light-inducible. We generated an integrable cassette containing the essential components of a blue-light reconstituted transcription factor. We chose to use a split transcription factor consisting of a DNA-binding domain fused to the naturally occurring *Arabidopsis* cryptochrome CRY2 photolyase domain (DBD-CRY2PHR) and the CIB1 protein fused to the VP16 activation domain (VP16-CIB1). In response to blue-light CRY2 undergoes a conformational change that allows it to bind CIB1, which recruits the VP16 activation domain to a promoter of interest containing binding sites for the selected DNA-binding domain (DBD) driving gene expression. We chose the DNA-binding domain of the Zif268 transcription factor (ZDBD), which is known to bind a 9-bp site (GCGTGGGCG) that has only 11 predicted binding sites in the *S. cerevisiae* genome^42^. Studies using the ZDBD on an estradiol inducible transcription factor have shown that artificial transcriptional activators using this DNA-binding domain in *S. cerevisiae* generate very little off-target gene expression activity^42,43^. When the Zif268 DNA-binding domain is fused to CRY2PHR, the resulting ZDBD-CRY2PHR/VP16-CIB1 transcription factor controls the expression of yeast genes under a pZF(BS) promoter containing GCGTGGGCG binding sites (BS) in a blue-light dependent manner^43^.

Stable integration of the ZDBD-CRY2PHR/VP16-CIB1 transcription factor is a more promising approach than maintenance of the optogenetic components on episomal plasmids, as expression from plasmids is known to be noisy and requires constant selection^44^. In order to integrate the ZDBD-CRY2PHR/VP16-CIB1 optogenetic machinery without loss of a marker, we used the heterologous URA3 from *Kluyveromyces lactis* (KlURA3) flanked by two direct repeats of the loxP sequence to allow for Cre recombinase mediated marker excision^45^. The components were cloned as indicated in **Figure 1A** using standard cloning techniques as described in the Methods. Homology arms on either side of the cassette allow for rapid integration at the HO locus, which is not required for growth and does not have an effect on growth rate^46,47^.

**Figure 1:**
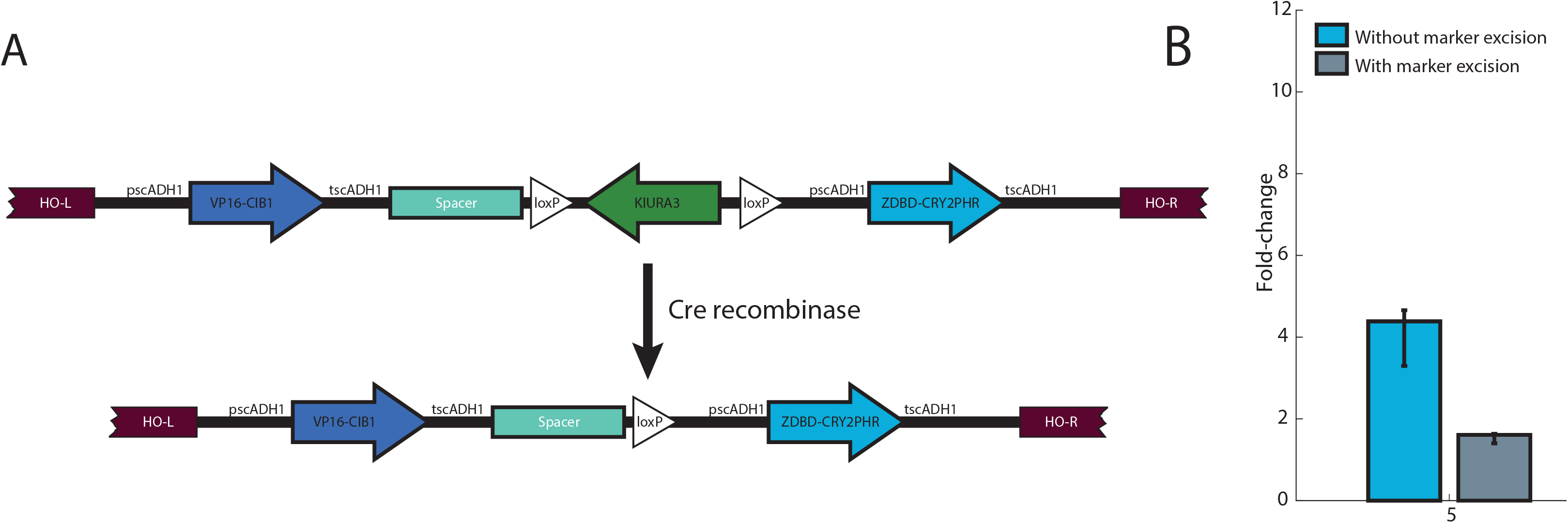
Vectors to integrate the blue light inducible split transcription factor (ZDBD-CRY2PHR/CIB1-VP16) into yeast with marker recovery. **(A)** The split TF vector inserts ZDBD-CRY2PHR and VP16-CIB1 at the HO locus under the expression of strong promoters with KIURA3 selection. Expression of Cre-recombinase and recombination of the loxP sites removes the KIURA3 marker, leaving it available for future strain manipulation. **(B)** Illumination of strains with ZDBD-CRY2PHR/CIB1-VP16 and a pZF(3BS)-yEVENUS reporter at 460nm (50μW/cm^2^) demonstrates that the ZDBD-CRY2PHR/CIB1-VP16 transcription factor drives gene expression from the pZF promoter in strains with or without recycling of the KIURA3 marker. However, removal of the KIURA3 marker does reduce expression from the pZF promoter approximately two-fold. Expression of yEVENUS was measured using imaging cytometry. Error bars are bootstrapped 95% confidence intervals for the mean expression level.

We also included spacer DNA of approximately the same length as KIURA3 as indicated in **Figure 1A** based on initial tests of the scheme which indicated that the spacing between the two open reading frames encoding the split transcription factor is important for optimal function of the optogenetic system (**Supplemental Figure 1**).

Integration of the ZDBD-CRY2PHR/VP16-CIB1 machinery at HO enables light dependent expression from the pZF(3BS) promoter in cultures grown in liquid and on solid media (**Figure 1B, Supplemental Figure 2**). Excision of the KIURA3 marker still results in some attenuation of gene expression in the marker-recycled strain (**Figure 1B**). We hypothesize that this is due to repression of ZDBD-CRY2PHR expression by the strongly expressed upstream VP16-CIB1 gene (**Figure 1A**). This could be due to terminator-promoter interactions as previously reported^48,49^. Previous work has shown that the ratio of CRY2PHR to CIB1 in the split transcription factor is important for maximal gene expression^43^ and it is possible that removing the KlURA3 marker changes the ratio to be slightly less favorable. We note that using higher light intensities (**Supplemental Figure 3A**) increases gene expression and that significant expression does not require a multi-copy reporter plasmid (**Supplemental Figure 3B**). In subsequent experiments, the reduced expression due to excision of the KIURA3 marker did not cause difficulties but we note that if maximal gene expression is required, constructs designed to optimize the dosage of VP16-CIB1 and ZDBD-CRY2PHR have been described^43^.

To allow specific genes in the yeast genome to be optogenetically controlled, we designed a cassette containing a pZF(BS) promoter (5’->3’) and the KanMX cassette (3’->5’) (**Supplemental Figure 4A**), which confers resistance to the antibiotic G418^50^. Replacing an endogenous promoter with this cassette in a strain containing the ZDBD-CRY2PHR/VP16-CIB1 split transcription factor puts expression of the gene of interest under blue-light control. We verified that in the dark, replacement of the native promoter with pZF(BS) effectively generates a deletion. Replacement of the HIS3 promoter with this cassette generates a histidine auxotroph (*his3*^*-*^) in the dark and the the ability to grow without histidine is recovered when grown in blue light in the presence of the ZDBD-CRY2PHR/VP16-CIB1 split transcription factor (**Supplemental Figure 5**). Gene expression from this promoter is rapid (7-fold gene expression in 2 hours) as assessed by pZF(BS)-yEVENUS (**Supplemental Figure 4B**). In combination, the cassettes containing an integrable light-responsive split transcription factor (ZDBD-CRY2PHR/VP16-CIB1) and a drug-selectable promoter cassette (KanMX4-pZF(BS) allow expression of a gene of interest to be put under light control in a variety of *S. cerevisiae* strains.

### Creation of a light-inducible invertase *S. cerevisiae* strain

We decided to take advantage of the well-understood invertase public goods system in budding yeast to generate yeast strains where cooperative intercellular interactions could be controlled by light (**Figure 2A**). In a yeast strain with the ZDBD-CRY2PHR/VP16-CIB1 optogenetic system stably integrated at HO we replaced the SUC2 promoter with pZF(3BS) (using the KanMX-pZF(3BS) cassette). Lawns of strains plated on YP-Sucrose were able to grow in blue-light, but not in the dark (**Supplemental Figure 6**) indicating that these strains induced SUC2 in a light-dependent manner, allowing the cells to produce invertase and utilize sucrose.

**Figure 2:**
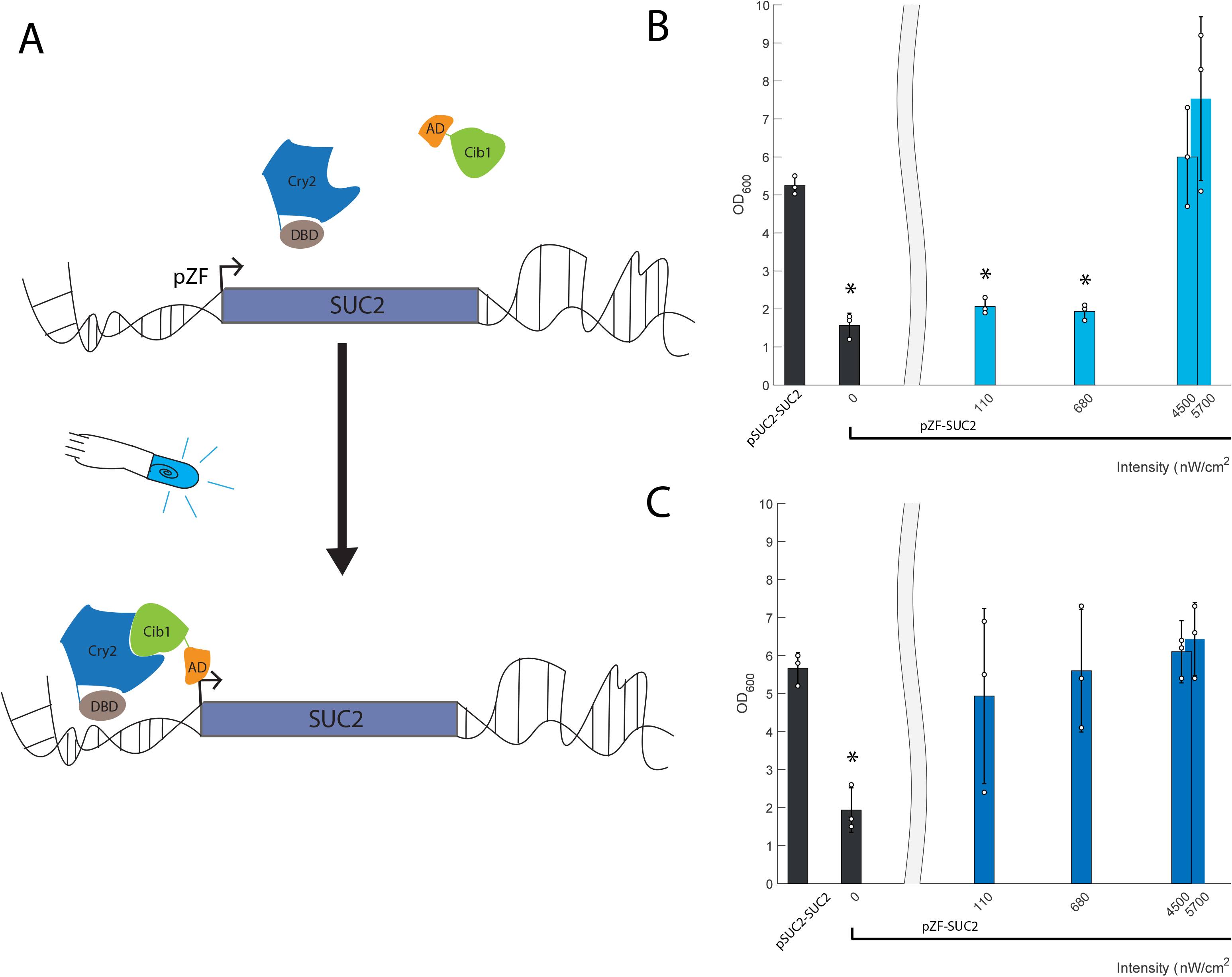
Characterization of a light inducible invertase strain. **(A)** In a yeast strain containing the ZDBD-CRY2PHR/VP16-CIB1 gene cassette the invertase endogenous promoter (pSUC2) was substituted with the orthogonal light inducible promoter (pZF) using the KanMX-pZF(3BS) cassette. In the dark, chimeric proteins ZDBD-CRY2PHR and VP16-CIB1 remain unbound and are inactive. Upon the addition of light CRY2 undergoes a conformational change that allows binding to CIB1 and recruits VP16-CIB1 to the promoter to drive transcription. The optogenetic strain pZF-SUC2 was exposed to a range of light intensities (0 nW/cm^2^-14 × 10^5^ nW/cm^2^) in YP-Sucrose media. **(B)** At 24 hours the wild-type strain (pSUC2-SUC2) shows robust growth, while the control (pZF-SUC2, 0 nW/cm^2^) does not. When provided a sufficient light dose, the pZF-SUC2 strain is able to recover wild-type growth in 24 hours (intensities >4000 nW/cm^2^) **(C)** After 48 hours, all pZF-SUC2 strains exposed to light catch up to the wild-type (pSUC2-SUC2) strain. Each bar represents three biological replicates and the individual data points are shown. (* denotes p<0.05, two-way ANOVA)

We further tested the ability of this strain to recover growth on YP-Sucrose in liquid cultures exposed to blue-light. We grew cultures over a range of light intensities (0 nW/cm^2^-14 × 10^5^ nW/cm^2^) and measured optical density after 24 and 48 hours of growth (**Figure 2B,C**). The parent strain (SUC2) quickly saturated at both 24 and 48 hours. In contrast, the pZF-SUC2 strain showed very little growth after both 24 and 48 hours of growth in the dark. Increasing intensity of blue light led to saturating optical densities at both 24 and 48 hours. Interestingly, at high light intensities (>4 µW/cm^2^) we reproducibly observed that pZF-SUC2 cultures reached a higher density than the wild-type pSUC2-SUC2 parent strain (yMM1146). It is possible that by decoupling production of invertase from the native regulation, the light-inducible strains overproduced invertase and are hence able to access more carbon from the sucrose. At low intensities of light (**Figure 2B**, 110 nW/cm^2^ and 680 nW/cm^2^) the culture did not show significant growth at 24 hours but by 48 hours was able to reach a wild-type level of saturation. This could be due to the known Allee effect^51,52,52–54^ (density-dependent growth) caused by the cooperative metabolism of sucrose by secreted invertase. At low intensities of light, low invertase production and secretion slows sucrose hydrolysis and population growth, delaying the point (relative to higher light intensity cultures) at which the population reaches a density that supports maximal growth rate.

We further tested the induction dynamics of our light-inducible strain over a several day growth experiment (**Figure 3**). The wild-type SUC2 strain quickly saturated after 20 hours of growth while the pZF-SUC2 strain had a delayed lag period, relative to the wild-type strain, which we interpret in light of the data in **Figure 2B** as needed time to accumulate invertase and glucose in the media after light induction. Subsequent to initiation of growth, the pZF-SUC2 strain showed very similar growth kinetics to the wild-type strain and quickly reached saturation. Again, the pZF-SUC2 reached a higher density than the wild-type strain at saturation, as we previously observed (**Figure 3**). Interestingly, the pZF-SUC2 strain also showed some growth in the dark, albeit after an extremely delayed lag period. We know from previous studies^43^ that the pZF promoter is not absolutely silent, and therefore we interpret this growth as being due to an extremely slow accumulation of functional invertase and hexose due to leakiness from the pZF promoter. We confirmed that our sampling method did not inadvertently expose cultures to unwanted light by demonstrating that the final densities of our time course samples did not show any significant difference relative to untouched endpoint samples (**Supplemental Figure 7**).

**Figure 3:**
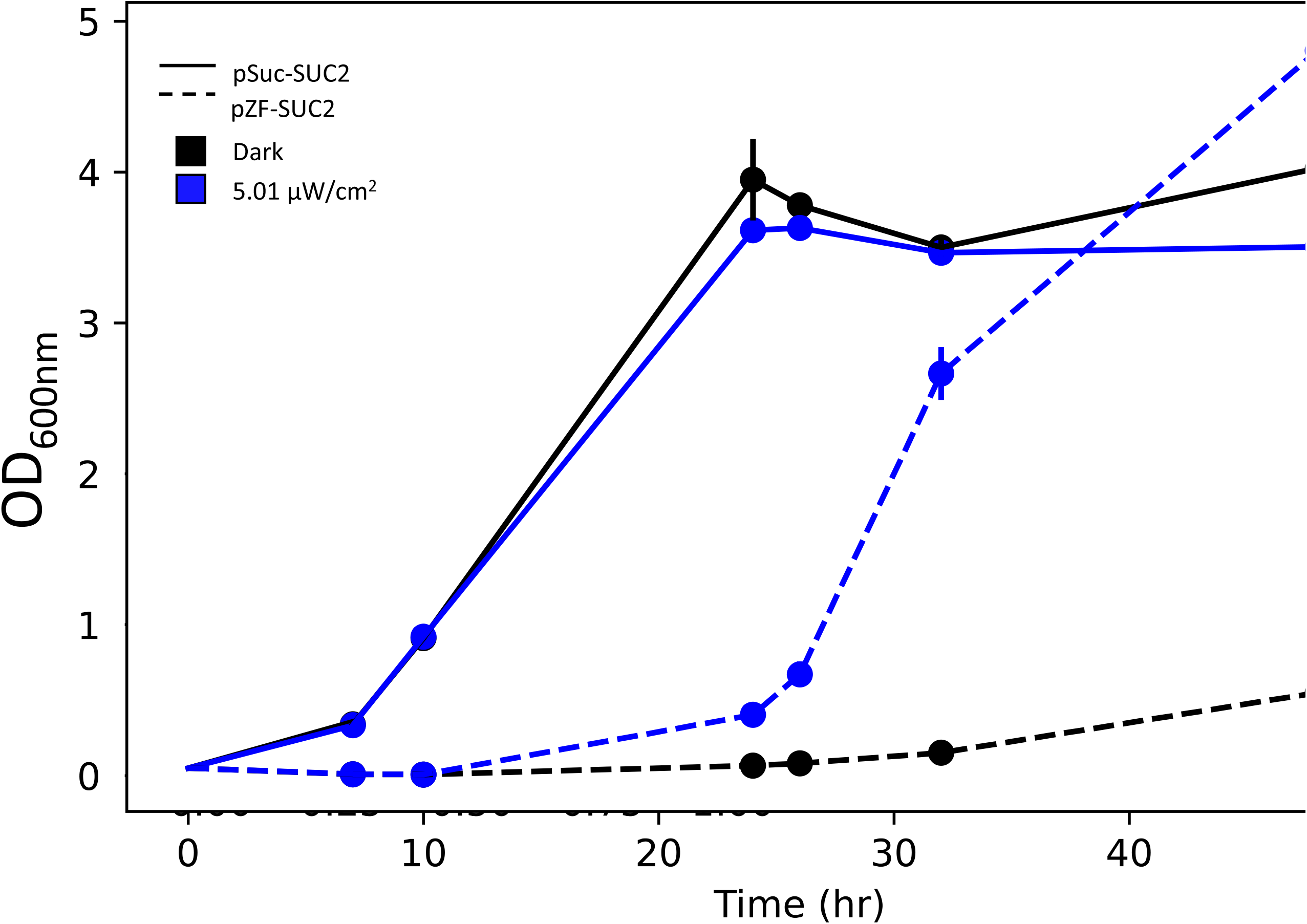
Light induction time course of light inducible strain (pZF-SUC2, dashed line) and wildtype strain (pSUC2-SUC2, solid line) at a light intensity of 5.01 µW/cm^2^ (blue) and no light (black) (n=2). Error bars are standard deviation. Error bars not visible are smaller than the marker. The optogenetically controlled strain displays a long lag in growth, which may be due to the time needed to accumulate invertase and break of sucrose to support growth after light induction.

### Light patterning allows for spatial control of producer populations

The experiments described above demonstrate that we can control invertase production, and therefore cooperativity, with blue light. However, these experiments were all done in well mixed populations, while microbial communities are generally highly structured 2D- or 3D-environments. Therefore, we wanted to test our ability to spatially control cooperativity in populations of *S. cerevisiae*.

Localized illumination of a regular grid of pZF-SUC2 strains arrayed onto an agar pad demonstrated that a small, spatially localized group of cooperators (**Supplemental Figure 8**) can support growth of a much larger number of cheaters in two-dimensional environments. This is expected due to diffusion of hexose. While the invertase enzyme is anchored to the plasma membrane, the fructose and glucose converted by the enzyme is free to diffuse and a relatively small fraction is captured by the cell that makes it^17^. To further validate this technique, we generated plates containing a lawn of pZF-SUC2 cells and illuminated a spot through a 6 mm pinhole (**Figure 4A)**. We found that after 4 days, the growth of very few cheaters was supported, with the majority of growth visible within the illuminated region. However, after 7 days of illumination, the cooperating cells supported a large growth of cheaters presumably because they were continuing to produce invertase and hydrolyze sucrose to hexose and the majority of hexose diffuses away from the illuminated cells (*i*.*e*. the producers).

**Figure 4:**
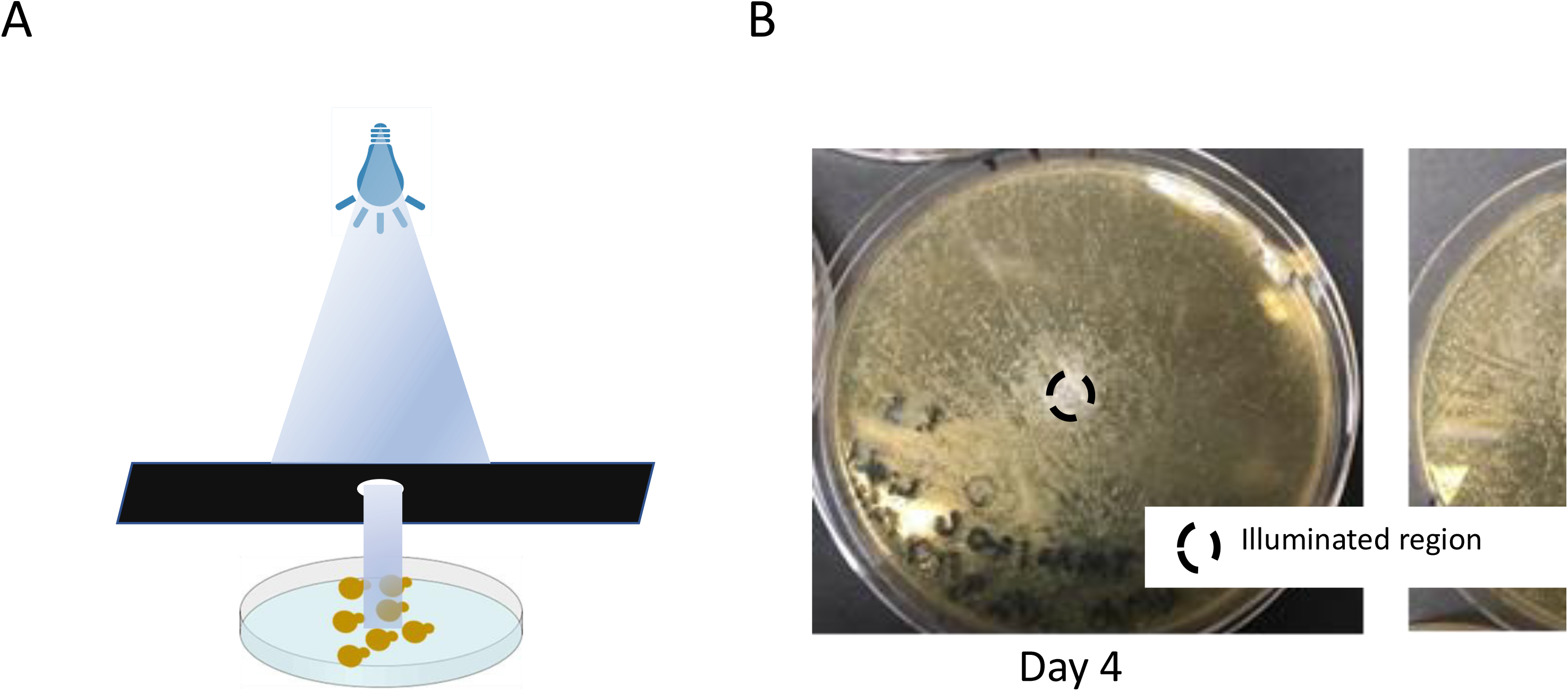
Controlled light results in patterned growth of a synthetic public goods community. (A) Spatial patterning of our public goods communities can be achieved by optogenetically controlling invertase expression in an illuminated area. A photomask limits the illuminated area on a petri plate resulting in patterned growth. (B) A representative image of the public goods community patterned on a standard petri plate. The black circle denotes the illuminated area of the plate. Growth was imaged on Day 4 and Day 7.

### Spatial patterning due to spatial segregation of cooperators and nutrient competition

In our pinhole experiment, we noticed a subtle ring effect (**Figure 4B**, Day 7), where the illuminated cooperators (at the center) grew well, surrounded by a ring of lesser growth, and finally more dense growth of cheaters at the periphery of the entire colony. This kind of ring-like pattern formation is predicted in reaction-diffusion systems where an activator and an inhibitor diffuse from a central source on different timescales^55–58^. In our system, the central cooperators are activating the growth of cheaters by producing hexose while simultaneously inhibiting the growth of cheaters by serving as a sink for other limiting growth factors (*i*.*e*. nutrients)^55–57^. Cheaters growing near the initially faster growing cooperators have access to hexose but are deprived of other nutrients, while cheaters at the periphery have more access to the limiting nutrient in the plate and eventually have access to hexose diffusing from the central cooperators, and therefore grow to a higher density.

In order to more fully explore this observation, we used auxotrophic strains to allow control of a limiting nutrient on the plates. Our wild-type pSUC2-SUC2 and opto-control pZF-SUC2 strains are leu2 auxotrophs allowing us to control leucine amino acid concentration in the plates to limit a nutrient. As a control, we generated a constitutive suc2Δ leu2Δ cheater strain. We spotted suc2Δ leu2Δ cheaters, pZF-SUC2, or pSUC2-SUC2 cells onto lawns of suc2Δ leu2Δ cheaters. Leucine concentrations in the plates were chosen to be 100% (0.1 mg/ml) or 50% (0.05 mg/ml) of the amount used in standard synthetic media^59^. As expected, spotting suc2Δ leu2Δ cheaters onto a lawn of suc2Δ leu2Δ cheaters does not allow for any growth in either light or dark **(Figure 5A)**. In contrast, both pZF-SUC2 or wild-type pSUC2-SUC2 cells spotted onto cheaters and grown in blue light allows for clear growth of the cooperators (either pZF-SUC2 or wild-type) surrounded by a zone where growth of the suc2Δ leu2Δ cheaters is inhibited and a larger ring of dense cheater growth **(Figure 5A, B)**. The growth inhibition zone is larger for wild-type cooperators than pZF-SUC2 cooperators **(Figure 5B, Supplemental Figure 9, 10)**. We interpret this to be due to more rapid induction of invertase and glucose production in the wild-type strains which allows the wild-type strain to more quickly reach a high density of cooperators allowing further cooperator growth (as also seen in **Figure 3**), and greater utilization and depletion of leucine. That leucine is the limiting nutrient is evidenced by lesser growth in both the wild-type and pZF-SUC2 strains at 50% leucine than at 100% leucine **(Supplemental Figure 9, 10)**.

**Figure 5:**
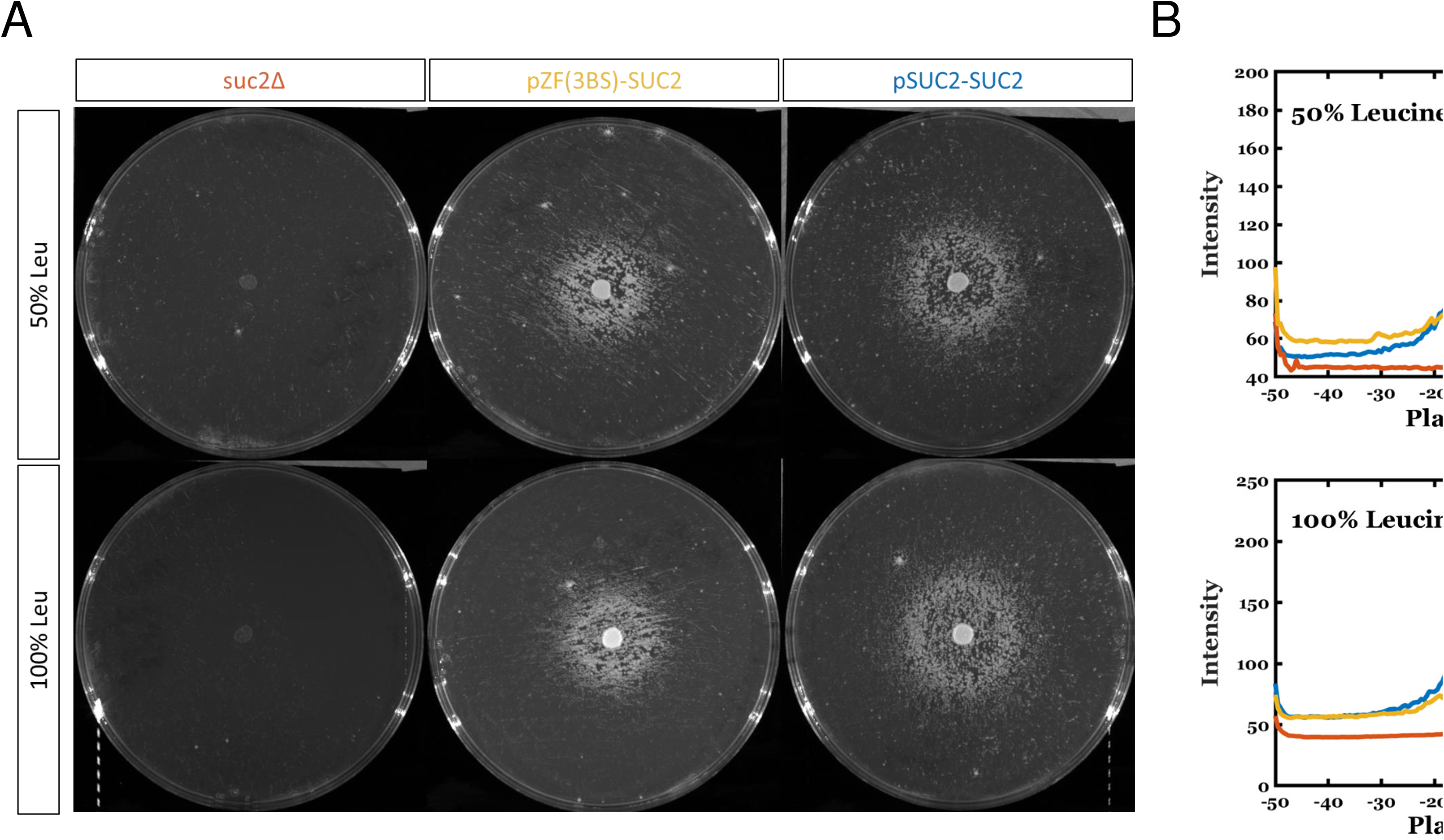
Spot patterning assay on nutrient limited SC-Sucrose plates. **(A)** Representative images of spot assay plates. The top row is composed of 50% leucine plates while the bottom row shows the 100% leucine condition. All plates were spread with the constitutive cheater strain, suc2Δ leu2Δ. From left to right the spotted strains are: suc2Δ leu2Δ, pZF-SUC2 and pSUC2-SUC2. **(B)** Plots showing an averaged radial intensity profile of the spotted plates across the diameter of the plate. At both concentrations of leucine, the pSUC2-SUC2 strain (blue) shows a larger inhibition of growth zone than the pZF-SUC2 (yellow) strain. In all cases, there is no growth of the suc2Δ leu2Δ(orange) strain.

## Discussion

This study develops and demonstrates the use of an optogenetic tool to control cooperation in a yeast microbial community. By making expression of invertase (encoded by the SUC2 gene) light-controllable we demonstrate temporal and spatial control of public goods production. We show that the timing of invertase expression is important, and delays in expression can significantly slow community growth. In addition, we show that localized cooperation can generate distinct patterning of cooperators and cheaters. Despite frequent investigation of *S. cerevisiae* invertase secretion as a model cooperative community, most models approximate invertase production as constant in time and space despite known native regulation in response to external factors such as nutrient concentration^60,61^. Optogenetic control of invertase will allow for further dissection of how regulation of this enzyme in space and time allows cooperators to coexist and compete with cheaters. While we have focused on the control of an intercellular interaction, the optogenetic constructs and strains generated in this study can be immediately used by other researchers to put any gene of interest under the control of blue light in *Saccharomyces cerevisiae*. The optogenetic system is orthogonal to native regulatory systems^43^ and could be easily modified to utilize additional markers or CRISPR technology for integration into a variety of yeast strains or species.

More generally, this study suggests that optogenetics will be a powerful tool for understanding how spatiotemporal regulation of cooperation, and other interactions, control microbial community structure and phenotype. Interactions in microbial communities are mediated by diffusible compounds, and numerous studies indicate that short-range interactions on the micron to millimeter scale are important for controlling community structure and phenotype^62–64^. Yet controlling the spatial arrangement of microbes on these length scales can be challenging. Microfluidic devices allow spatial segregation of microbes at different length scales but require sophisticated engineering and specialized equipment^65–68^. In addition, it is more challenging to define three-dimensional structure using a microfluidic device and collection of the community for subsequent downstream analysis (*e*.*g*. gene expression) can be difficult. Bioprinting is a burgeoning technique which holds promise for building complex, three-dimensional microbial communities with defined spatial structure^65,69–71^. However, bioprinting does not easily allow intercellular and interspecies interactions to be modulated in time. Optogenetics has the potential to be integrated with or to supersede these existing technologies for fine spatiotemporal control of community interactions. Scanning and parallel light-targeting methods can be combined with one and multi-photon excitation to precisely localize light in both two and three dimensions as well as in time. In addition, existing illumination techniques can be combined with amenable animal models, such as *C. elegans*^72^, to allow unprecedented *in vivo* dissection of the importance of intercellular interactions and their regulation in the establishment and phenotype of microbial communities. To extend the techniques described in this article to mixed-kingdom communities, optogenetic systems developed for bacteria^80,81^ could be utilized. Indeed in *Sinorhizobium meliloti*, a nitrogen fixing soil bacterium, the blue-light sensitive transcription factor EL222 was recently used to control production of the public good exopolysaccharide enabling manipulation of biofilm formation^41^. Hybrid optochemical approaches also hold promise for repurposing existing inducible systems, as a recent study showed that photocaged IPTG could be used to control coculture interactions in the bacterium *Corynebacterium glutamicum*^39^.

Finally, in addition to providing a path towards understanding how intercellular interactions regulate naturally occurring microbial communities, optogenetic tools have important implications for engineering synthetic microbial consortia. Engineered consortia are of great interest in biotechnology because they can perform more complicated functions than single-species or single-strain communities^73^. However, maintaining the appropriate ratio of different consortia members represents a challenge and would benefit from dynamic control modalities. Control mechanisms for cocultures via interspecies interactions (such as quorum sensing and metabolite exchange) have been described^74–76^ and dynamic control of these interactions using optogenetics and predictive control strategies^36,77^ could enable community maintenance and optimization. Similar optogenetic approaches in monocultures have already enabled significant gains in bioproduction^35,78,79^. In addition, the spatial control provided by light could allow the formation of sophisticated living biomaterials. Co-cultures of *Saccharomyces cerevisiae* and the cellulose-producing *Komagataeibacter rhaeticus* bacteria are mediated by yeast invertase production and capable of producing functionalized cellulose biomaterials. Optogenetic control of *Saccharomyces cerevisiae* invertase production could allow for sophisticated control of these living materials as well as patterning, as demonstrated in this work for the simple case of localized producers. In addition, as demonstrated by the grid experiment (**Supplemental Figure 8**) a small number of producers can support a much larger population, indicating that in living materials it may be possible to have a relatively small population responsible for the metabolic burden of consortia growth while other members can focus on additional functionality.

## Materials and Methods

### Yeast Strains and Culture Methods

Yeast strains used in this study are shown in **Supplemental Table S1**. Yeast transformation was accomplished using standard lithium-acetate transformation^82^. For integrating plasmids, the integration was validated using either colony PCR or, when colony PCR proved difficult, by PCR of genomic DNA. Genomic DNA was extracted using the Bust n’ Grab protocol^83^. Primers used for validating integrations are listed in **Supplemental Table 2**. All transformants were checked for the petite phenotype by growth on YEP-glycerol (1% w/v Bacto-yeast extract-BD Biosciences 212750, 2% w/v Bacto-peptone-BD Biosciences 211677, 3% [v/v] glycerol-Fisher Bioreagents BP229-1, 2% w/v Bacto-agar-BD Biosciences #214030)[22]. Only strains deemed respiration competent by growth on YEP-glycerol were used for subsequent analysis. Details of individual strain construction are described in the **Supplemental Material**.

Yeast cultures were grown in either yeast peptone (YP) media (10 g/L Bacto yeast extract, 20g/L Bacto peptone for solid media + 20g/L of Bacto agar) or Synthetic Complete (SC) media (6.7 g/L Yeast Nitrogen Base without amino acids-DOT Scientific, 1% v/v KS amino acid supplement without appropriate amino acids). The carbon source supplied was either dextrose (D) or sucrose (SUC) at 2% v/v concentration. As needed, episomal plasmids were maintained by growing yeast in SC media lacking the appropriate amino acids required for plasmid selection. For light induction experiments followed by fluorescence assays (flow cytometry or microscopy) yeast were always grown in Synthetic Complete media^59^.

### Bacterial strains and growth media

*Escherichia coli* strain DH5α was used for all transformation and plasmid maintenance in this study. *E. coli* were made chemically competent following either the Inoue method^84^ or using the Zymo Research Mix & Go! Protocol (Zymo Research T3002). *E. coli* were grown on LB agar (10% w/v Bacto-Tryptone, 5% w/v Bacto Yeast Extract, 5% w/v NaCl, 15% w/v Bacto Agar) or LB liquid media (10% w/v Bacto-Tryptone, 5% w/v Bacto Yeast Extract, 5% w/v NaCl). Appropriate antibiotics were used to select for and maintain plasmids. Antibiotic concentrations used in this study were as follows: LB+CARB agar 100 μg/mL carbenicillin, LB+CARB liquid media 50 μg/mL carbenicillin, 25 µg/ml chloramphenicol, 50µg/ml kanamycin. Plasmids were prepared using the Qiagen bacterial miniprep kit (Qiagen #27104).

### Plasmid Construction

Construction of plasmids used throughout this study was accomplished using a combination of methods including Yeast Recombinational Cloning^85^ and standard restriction enzyme based cloning. Details of individual plasmid construction are described in the **Supplemental Materials**.

### Generation of optogenetic invertase strain

In order to make an integrable version of the SV40NLS-VP16-CIB1 loxP-KlURA3-loxP SV40NLS-ZIF268DBDCRY2PHR cassette, this cassette was cut from pMM364 using XbaI/PacI and ligated into pMM327. This plasmid was linearized using AatII and transformed into yMM1146 (Matα trp1Δ63 leu2Δ1 ura3-52) to generate yMM1367 (Matα trp1Δ63 leu2Δ1 ura3-52 HO::SV40NLS-VP16-CIB1 loxP-KlURA3-loxP SV40NLS-Zif268DBD-CRY2PHR). The KIURA3 marker was excised from this strain using Cre-mediated recombination as described below to generate yMM1390 (Matα trp1Δ63 leu2Δ1 ura3-52 HO::SV40NLS-VP16-CIB1 loxP SV40NLS-Zif268DBD-CRY2PHR). In order to make the expression of invertase light-inducible, the pZF(3BS) promoter replaced the native pSUC2 promoter by amplifying KanMX-rev-pZF(3BS) with oMM768/769 from pMM353, transforming yMM1390 and selecting for G418 resistant colonies. These colonies were further checked by colony PCR and sequencing and for the inability to grow on YP-Sucrose in the dark and became strain yMM1406.

### Recycling of loxP-flanked markers

The Cre-loxP system was used to recycle the KIURA3 marker flanked by loxP recombination sites (loxP-KIURA3-loxP). Cre-mediated recombination was accomplished by adapting the CRE recombinase-mediated excision protocol from Carter and Delneri (2010)^86^. The strain yMM1367 (Matα trp1Δ63 leu2Δ1 ura3-52 HO::SV40NLS-VP16-CIB1 loxP-KlURA3-loxP SV40NLS-Zif268DBD-CRY2PHR) was transformed with 0.25-0.5µg of pMM296 (pSH65, pGAL1-CRE Bleo^R^). These transformants were plated onto YPD and then replica plated onto selective media (YPD +10µg/ml phleomycin (InvivoGen)) after overnight growth. To express CRE and induce recombination phleomycin resistant colonies were selected and grown overnight in 3ml of YP-Raffinose (1% w/v yeast extract (BD Biosciences), 2% w/v Bacto-peptone (BD Biosciences), and 2% w/v raffinose (Becton Dickinson 217410)). The following day, cells were harvested by centrifuging at 3750 rpm for 5 minutes, washed in sterile miliQ water, and resuspended in 10ml of YP-Galactose (1% w/v yeast extract (BD Biosciences), 2% w/v Bacto-peptone (BD Biosciences), 2% w/v galactose (BD Biosciences 216310)) at an OD_600_ of 0.3. These cultures were incubated at 30°C with shaking for 2-3 hours. This culture was then diluted and plated on YPD and then replica plated onto SC-5FOA (25% w/v g Bacto-Agar, 6.72% w/v YNB, 1% v/v mL 20x KS supplement without URA, 2% v/v glucose, 10 mL 5-Fluoroorotic Acid (Zymo Research), 50 mg uracil (MP Biomedicals 103204). 5FOA resistant colonies were checked for excision of the KIURA3 marker using colony PCR. Transformants with KlURA3 excised were grown in liquid YPD to saturation twice and then plated on YPD for ∼100 colonies per plate. These were replica plated onto YPD + 10µg/ml phleomycin. Phleomycin sensitive colonies (colonies that had lost the plasmid pMM296) were reconfirmed by colony PCR to have loxed out KlURA3. This generated yMM1390 (Matα trp1Δ63 leu2Δ1 ura3-52 HO::SV40NLS-VP16-CIB1 loxP SV40NLS-Zif268DBD-CRY2PHR).

### Blue light induction of yeast cultures in liquid media

For blue light induction experiments in liquid media light was applied in one of three ways: 1) <u>Peripheral Illumination:</u> Cultures were grown in glass culture tubes on the outside lane of a roller drum at room temperature. Control (dark) samples were put in test tubes wrapped in foil on the inner lane of the roller drum. Three LEDs outputting 460nm blue light (Sparkfun #COM-08718) were placed at the three, nine, and twelve o’clock positions of the roller drum and turned on at T=0 (∼3 mW/cm^2^ at the LED; ∼25 *μ*W/cm^2^ at the sample) as described previously^35,43^. 2) <u>Bottom Illumination:</u> Cultures growing in glass tubes in a roller drum were directly illuminated from the bottom of the glass culture tube by LEDs mounted into the roller drum. The circuit was composed of three LEDs per tube (Sparkfun, #COM-09662), resistors of varying strength (Sparkfun, #COM-10969) and a 12 V power supply (LED supply, #12V-WM-xxA). 3) <u>Light Plate Apparatus:</u> Cultures were grown in 24 well plates (Arctic White, #AWLS-303008) and placed on a Light Plate Apparatus (LPA)^87^. The Light Plate Apparatus (LPA) is a published optogenetic tool which provides programmed illumination to each well of a 24-well plate. We assembled our LPA as described in^87^ and calibrated as previously described^88,89^.

For all illumination methods, response was assessed by flow (traditional or imaging) cytometry as described below or measurement of optical density. The light output of all light sources was measured and validated with a standard photodiode power sensor (Thorlabs, #S120VC) and power meter (Thorlabs, #PM100D) as previously described^43,88,89^.

### Blue light induction of drug resistance or restoration of histidine auxotrophy

To assess blue-light induction of drug resistance from a pZF(3BS)-NatMX plasmid, strain yMM1355 (Matα trp1Δ63 leu2Δ1 ura3-52 HO::GAL4AD-CIB1 loxP-KLURA3-loxP FLAG(3X)-SV40NLS-Zif268DBD - CRYPHR) was transformed with the pZF(3BS)-NatMX plasmid (pMM369) or an empty vector control (pMM6). Growth was assessed in the presence of clonNat (nourseothricin, 50μg/ml, YPD plates) in either 450nm blue light (50 *µ*W/cm^2^) or the dark by frogging saturated cultures at 1:10 dilution series onto the appropriate plates and growing for 2 days at 30°C in either light or dark. To assess recovery of histidine auxotrophy, strain yMM1295 was transformed with appropriate combinations of pMM284 (ZDBD-CRY2), pMM159 (GAL4AD-CIB1), pMM6 (ø), and pMM7 (ø) and saturated overnight cultures were frogged at 1:10 dilutions onto either SC or SC-Leu-Trp-His. Plates were grown at 30°C under 460 nm blue light or in total darkness. All strains grew on fully supplemented SC in either the light or dark (data not shown). Results for SC-Leu-Trp-His with and without light are shown in **Supplemental Figure 5**.

### Growth in sucrose media at different blue light intensities

Biological replicates were picked from a single colony on a YPD plate and transferred to a glass culture tube containing 5 mls of YPD media and grown to saturation overnight in the dark. The saturated culture (1.5ml) was pelleted by centrifuge (Eppendorf, #EP5401000137), washed twice with sterile water and resuspended in sterile MilliQ water to wash out residual media. These concentrated cells were then diluted at 1:100 into 5mls of SC-SUC to OD_600_∼0.16 and placed in a roller drum with the corresponding light dose, or wrapped in aluminum foil for the no light control. The cultures were allowed to grow for a total of 48 hours with 100uL of sample taken every 24 hours in order to measure the OD600 of the culture using a spectrophotometer (Fisher Scientific,#14-385-445).

### Time course of growth with light induction in sucrose media

A single yeast colony of yMM1406 and yMM1146 were inoculated into 5mls of YPD and grown overnight to saturation. Of these cultures, 1ml was pelleted and the pellet was washed three times with sterile water to washout residual media. These cultures were then resuspended and diluted in SC-sucrose media to an OD_600_ of 0.05. Each culture was divided into 12 wells of a 24-well plate (2ml of the diluted cultures) with a glass bead (Fisher Scientific, #11-312B 4mm) to increase aeration and the plates were covered with a breathable sealing membrane (USA Scientific #9123-6100) to reduce evaporation. Three light doses were programmed into the LPA with the arbitrary IRIS units of 0, 250, and 500. These correspond to 0*μ*W/cm^2^, 2.32 *μ*W/cm^2^, and 5.01 *μ*W/cm^2^, respectively. This resulted in a set of four wells for each strain at each light condition. Two of these wells were sampled at each timepoint, while two were left untouched until the final endpoint measurement to verify that intermediate manipulation of the plate did not inadvertently expose cultures to light. At each time point 100uL of the culture was removed to measure the optical density of the culture, the sealing membrane was replaced, and the plate was returned to the incubator. Optical densities outside of the previously determined linear range of our spectrophotometer were diluted to be in the linear rang at a ratio of 1:10 or 1:100 as needed. The experiment run-time was 54 hours.

### Blue light patterning

#### Patterning of yeast plates with blue light

Yeast strain yMM1406 (optogenetic producer) was inoculated into a 5mL test tube of YPD and grown overnight to saturation.. The next day the culture was pelleted by microcentrifuge (Eppendorf, #EP5401000137) at 3000G for 2 minutes, resuspended in sterile water to wash out residual media from the cell pellet, this process was repeated twice. The final OD 600 of the yeast cells was measured at 0.119 using a spectrophotometer (Fisher Scientific,#14-385-445). 200uL of the cell suspension was plated on YP-SUC plates and spread throughout the plate using glass beads (Fisher Scientific, #11-312B 4mm). The plates were wrapped in sterile, construction paper photomasks with one, 6 mm hole placed at the center of the plate or two 6mm holes placed at two different distances from each other and aluminum foil backing to prevent light contamination and control plates of no photomask (full-light at ∼57 µW/cm^2^) or complete photomask (no-light). The plates were placed under a blue light LED array and allowed to grow at room temperature for a week (until growth appeared to stagnate). Pictures of the plates were taken on day 4 and day 7 with a 28mm, 12 mega-pixel camera (iPhone 7).

#### Frogger plate patterning

Yeast strain yMM1406 (optogenetic producer) was inoculated into a 5mL test tube of YP-D to grow overnight, the culture was set back to an OD600nm of OD 0.219 and grown for a few hours until an OD600nm of 0.538 was reached. The culture was pelleted in a microcentrifuge (Eppendorf, #EP5401000137) at 3000G for 2 minutes and washed 3x with sterile water to wash away residual media. The culture was diluted to an OD600nm of 0.079 measured with a spectrophotometer (Fisher Scientific,#14-385-445) and a frogger tool (Dan-Kar corp, #MC48) was used to stamp a large culture plate (Corning, #431111) of YP-D agar, a photomask was placed over the bottom of the plate and only a small section of the plate (2cm^2^) was exposed to light at an intensity ∼145 µW/cm^2^ under a blue light LED array (HQRP New Square 12-inch Grow Light Blue LED 14W). The light source, and plate were placed in 30°C incubator. A lightbox (Amazon, #ME456 A4 LED Light Box) was used to illuminate the plate from the bottom and camera (gel box camera and hood) was used to image the plate on day 4.

#### Spot assay with blue light

Yeast strains yMM1146(wildtype producer), yMM1456 (non-producer) and yMM1406 (optogenetic producer) were inoculated into a 5mL test tube of YP-D to grow overnight. Cells were pelleted using a microcentrifuge (Eppendorf, #EP5401000137) at 3000G for 2 minutes and washed with YP-SUC to remove residual media containing dextrose, this was repeated 3 times. All cultures were diluted to an OD600nm of 0.04 measured with a spectrophotometer (Fisher Scientific,#14-385-445) before plating onto solid YP-SUC plates (Fisher Scientific, #BP94S01). Onto the lawns of suc2Δ leu2Δ cheaters growing on either 0.1mg/mL or 0.05mg/mL leucine we spotted 5µL of either suc2Δ leu2Δ cheaters, pZF-SUC2, or wild-type cells. Plates contained leucine concentrations of either 100% (0.1 mg/ml) or 50% (0.05 mg/ml) of the amount used in standard synthetic complete media^59^. All plates were spread with 150uL of the yMM1456 (suc2Δ leu2Δ) strain with glass beads (Fisher Scientific, #11-312B 4mm), the beads were removed and the plate allowed to dry for 10 minutes. Then, a 5uL drop of either yMM1146,1406, or 1456 was applied to the center of a petri dish and left face-up to dry for another 10 minutes. The plates were then placed upside down in a 30°C incubator in a single layer under a blue LED light source at an intensity 145 µW/cm^2^ (HQRP New Square 12-inch Grow Light Blue LED 14W) for 7 days. On the seventh day the pictures were imaged with ChemiDoc imaging system (BioRad, #12003154) an exposure of 0.06 seconds in the brightfield setting and analyzed using an ImageJ plug-in Clockscan (reference).

### Quantification of Plate Growth

#### Radial intensity traces of patterned plates using Clockscan

The patterned plates were analyzed using a published ImageJ plug-in, Clock Scan^90^ which outputs averaged radial intensity values for the image.

#### Identifying pattern features using custom MATLAB Script

We quantify the growth of yeast on a plate from images using a custom MATLAB script that examines intensity versus radius along angular slices through the center of the plate and identifies the bounds of features such as valleys and rings. Because it’s hard to accurately identify these features from individual angular slices or the single, composite intensity profile given by a clockscan^90^, we use a bootstrap-based approach to repeatedly identify potential features from randomly selected sets of angular slices and select the most frequently identified potential features as true features.

This starts by roughly identifying the central yeast spot using MATLAB’s circle finder and cropping the image around this spot. A polar transformation is then applied to the cropped image to create a polar image where each column of pixels corresponds to an angular slice through the plate. These angular slices are then sampled with replacement to construct a composite image. An intensity profile is generated from each composite image by taking the median intensity value at each radius. The intensity profile is filtered to remove noise and features are identified from the resulting signal. For example, potential valley bounds are identified as the locations where the derivative of the filtered intensity profile is at its maximum and minimum. This process is repeated for hundreds of composite images to create distributions of potential features. True features are then selected as the mode of these distributions. Using MATLAB’s circle find to identify the outer edges of the plate, which we know to be 100 mm across, we then convert the feature measurements to physical units. Code is available upon request.

### Flow Cytometry

Gene expression in response to blue light was assayed using fluorescent reporters and either traditional or imaging flow cytometry. Traditional flow cytometry was performed on a BD Biosciences LSR II Flow Cytometer (488nm laser and 505LP dichroic filter). The flow cytometry data was then analyzed using custom Matlab scripts. Imaging cytometry was done with the ImageStream MarkII and analysis completed using the IDEAS software or custom Matlab/ImageJ scripts modified from those described in [29].

All samples from culture tubes were prepared by diluting yeast cell culture (250-500µl) into 800µl of ice-cold PBS + 0.1% Tween-20. Samples were kept on ice or at 4°C until being analyzed. Samples from the Light Plate Apparatus (LPA) were taken by transferring 50µl of culture from each well of the LPA to a well a 96-well plate containing 150 μL of PBS+0.1% Tween-20. Samples run on the LPA were measured without sonication. Samples grown in glass culture tubes were sonicated with 10 bursts of 0.5 seconds each once diluted in PBS and prior to flow cytometry.

## Acknowledgements

This work was financially supported by the National Institutes of Health [R35GM128873] and a Lewis-Sigler Fellowship from Princeton University (M.N.M.). Flow cytometry was enabled by the University of Wisconsin Carbone Cancer Center Support Grant P30 CA014520 as well as the Princeton Flow Cytometry Resource Facility. Megan Nicole McClean, Ph.D holds a Career Award at the Scientific Interface from the Burroughs Wellcome Fund. Neydis Moreno Morales was supported by a short-term Genomic Sciences Training Program NHGRI Training Grant (5T32HG002760) and the Science and Medicine Graduate Research Scholars (SciMed GRS) program at the University of Wisconsin-Madison.

N.M.M., C.S., and M. P. designed experiments and performed experiments. N.M.M., C.S., and M.P. performed molecular cloning and strain construction. K.S. wrote image analysis software. N.M.M. and M.N.M. analyzed data and wrote the paper. N.M.M., C.S., M.P., K.S. and M.N.M. edited and approved the manuscript. M.N.M. supervised the project.

## Figure Captions

**Supplemental Figure 1:**
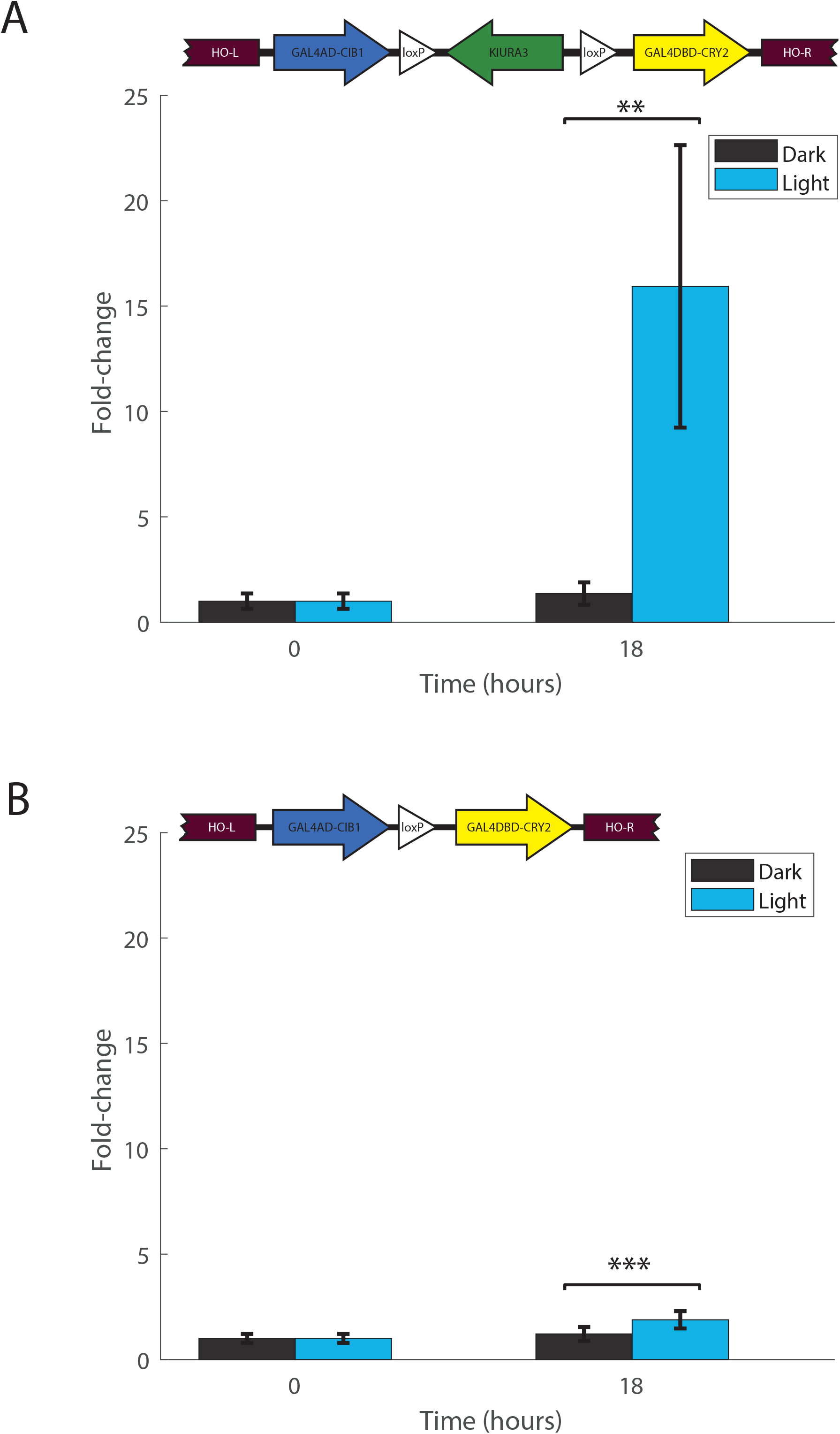
(**A**) Yeast cells carrying an integrated copy of a light activatable split GAL4 transcription factor (GAL4DBD-CRY2/VP16-CIB1) integrated at the HO locus without subsequent excision of the KlURA3 marker (inset) and a reporter plasmid (pGAL1-yEVENUS) were exposed to ∼ 25 µW/cm^2^ light for 18 hours. This resulted in an 15-fold induction in yEVENUS (p-value <0.005; Welch’s t-test). (**B**) In contrast, when Cre-recombinase is used to excise the KIURA3 marker (inset), only 2-fold induction in yEVENUS occurs after 18 hours of growth in ∼ 25 µW/cm^2^ blue light (p-value<0.0005; Welch’s t-test). Error bars are the standard error of the mean from 3 replicates.

**Supplemental Figure 2:**
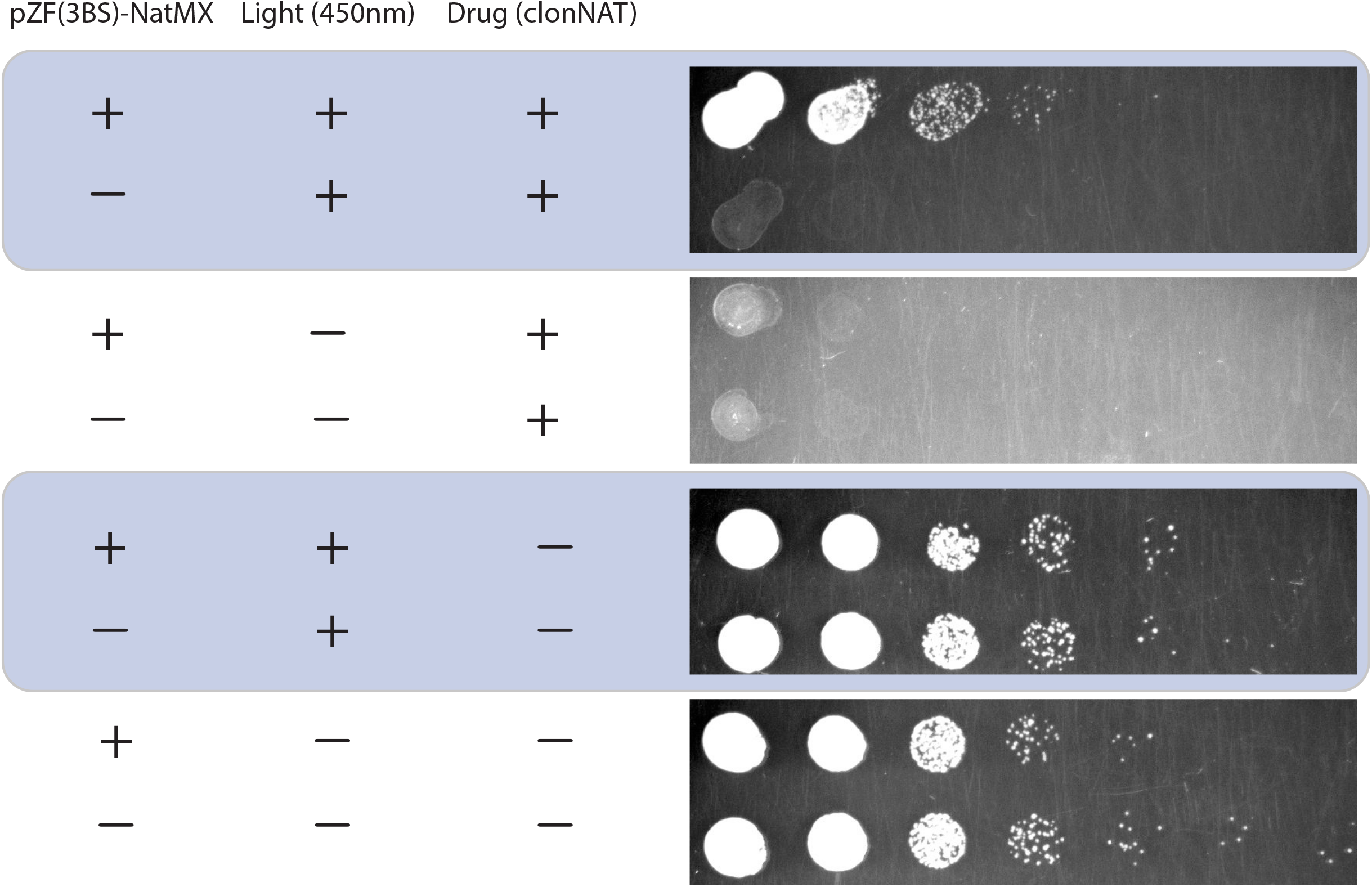
Yeast strains with the integrated optogenetic system and either a plasmid containing pZF(3BS)-NatMX or the empty vector control were grown in the presence of clonNAT (nourseothricin, 50 µg/ml) in either 450nm blue light (50 µW/cm^2^) or the dark (1:10 dilution series). Induction of the NatMX resistance marker (nat1 gene) by blue-light confers the ability to grow on nourseothricin.

**Supplemental Figure 3:**
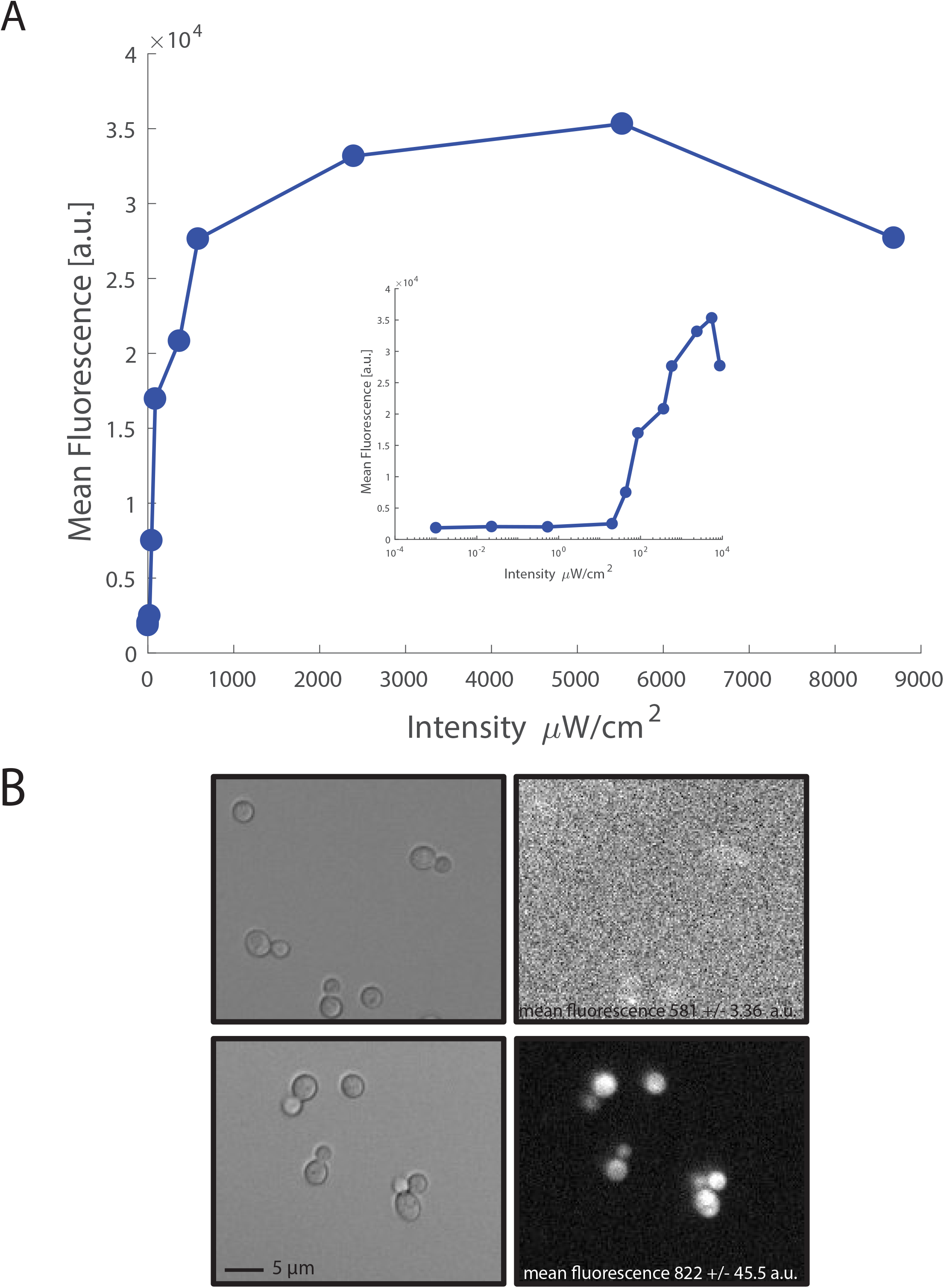
Expression from the pZF(3BS) promoter in blue light (**A**) The marker recycled strain (HO::SV40NLS-VP16-CIB1 loxP SV40NLS-Zif268DBD-CRY2PHR [pZF(3BS)-yEVENUS]) was grown for 18 hours in 460nm blue light over a range of intensities (0 *μ*W/cm^2^-8 mW/cm^2^). At least 20 *μ*W/cm^2^ of light is required to induce significant expression from the pZF(3BS) promoter (see inset) under these conditions. Error bars are the mean absolute deviation. (**B**) Integration of a pZF(3BS)-mRUBY2 reporter at the LEU2 locus in the unrecycled strain (HO::SV40NLS-VP16-CIB1 loxP-KlURA3-loxP SV40NLS-Zif268DBD-CRY2PHR) confers blue-light dependent expression on mRUBY2 (50 *μ*W/cm^2^ 460nm blue light, 18 hours, bottom row. Dark control-top row).

**Supplemental Figure 4:**
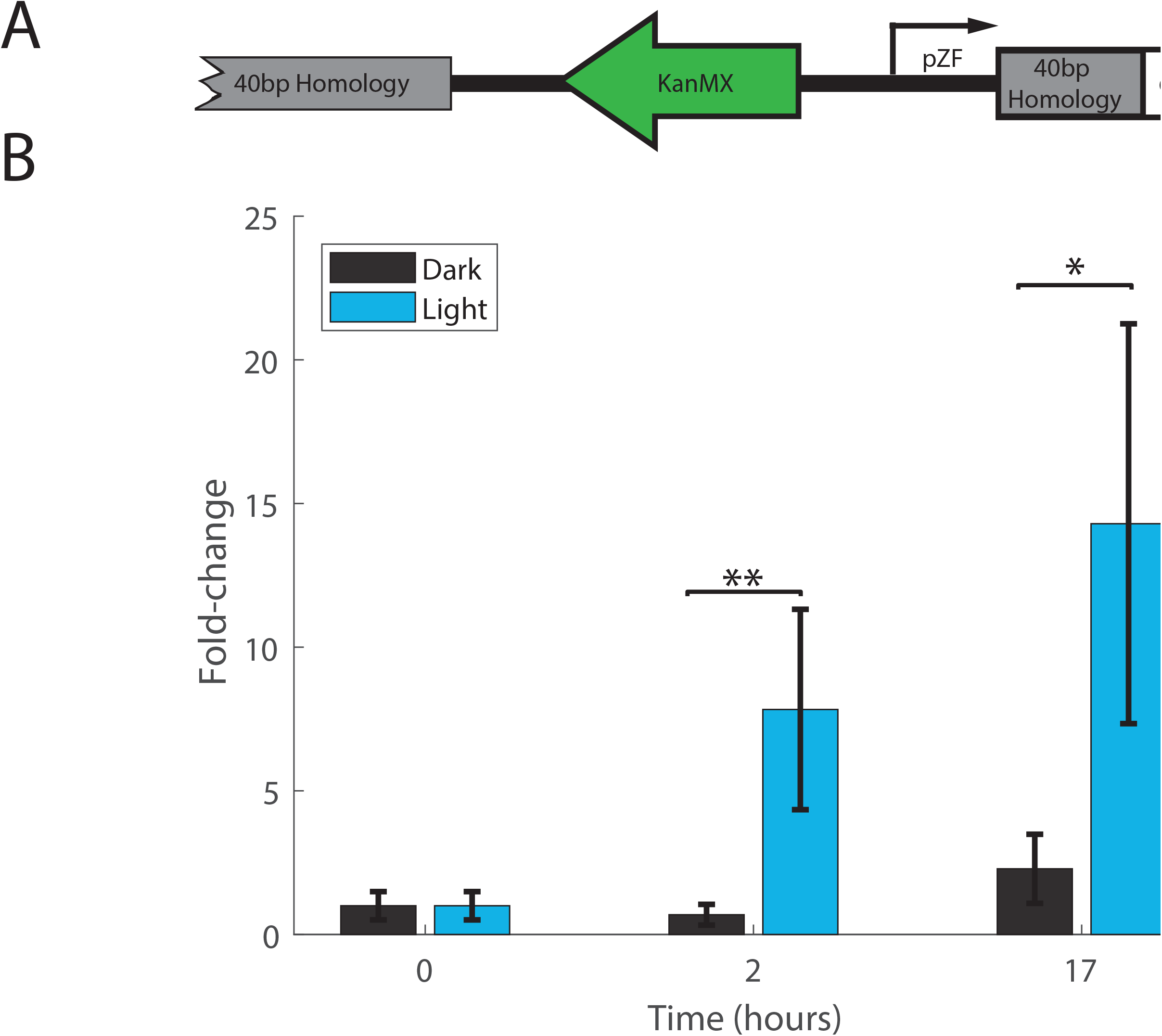
**(A)** Integration of the pZF promoter in front of a gene of interest using homologous recombination in yeast confers light-regulation. **(B)** Expression of a yEVENUS reporter from the pZF promoter under 25 *μ*W/cm^2^ 460nm illumination shown as fold-change relative to the T=0 dark sample. There is significant induction as early as T=2hrs of blue light (**p-value<0.005, * p-value<0.05; Welch’s t-test). Error bars indicate standard error of the mean fold-change.

**Supplemental Figure 5:**
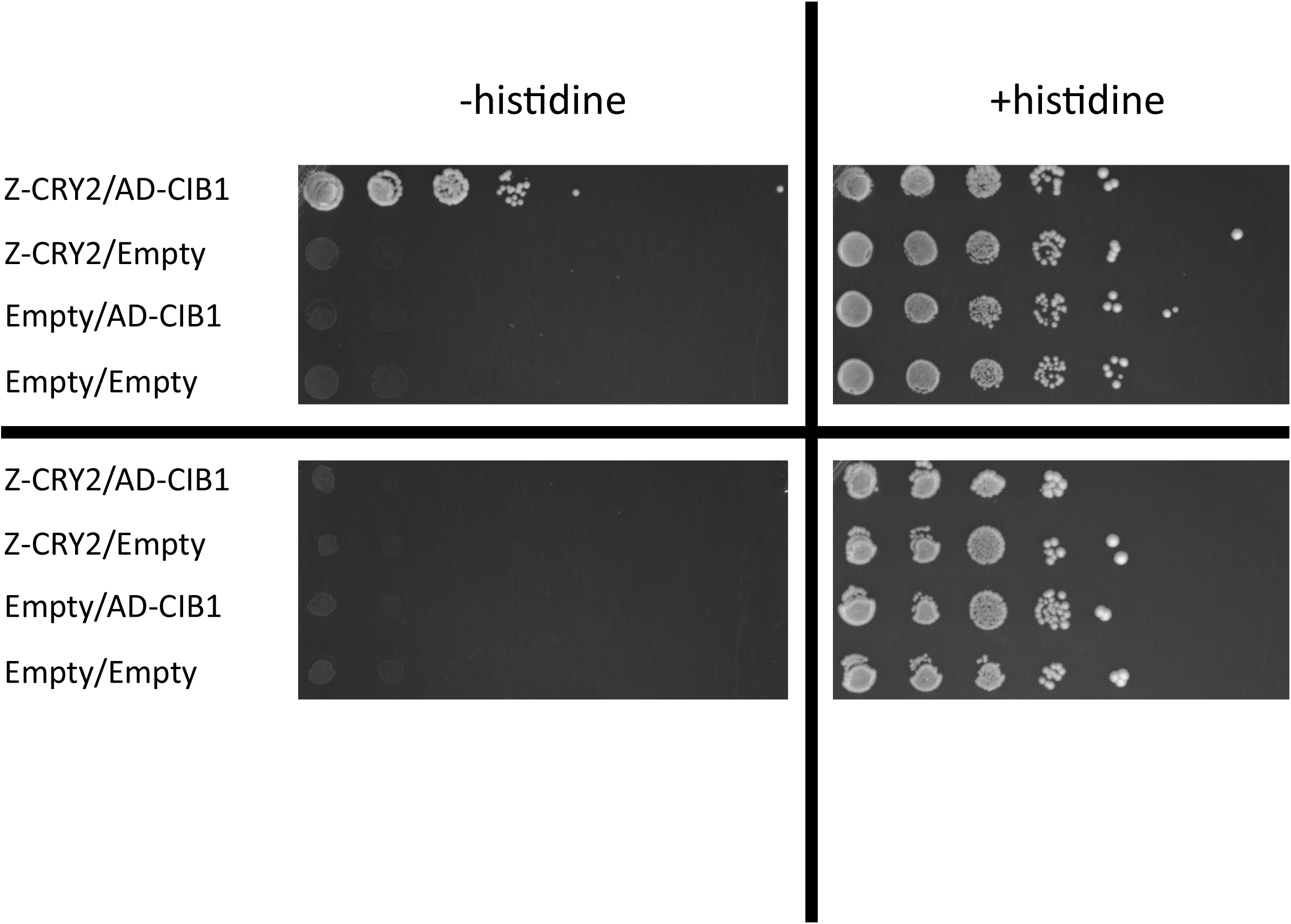
A yeast strain expressing imidazoleglycerol-phosphate dehydratase under the pZF promoter (pZF-HIS3) transformed with appropriate combinations of ZDBD-CRY2 (pMM284), AD-CIB1(pMM159) and/or empty vector controls (pMM6, pMM7) was frogged onto media without or with histidine at 1:10 dilutions either in the presence or absence of 57 µW/cm^2^ 460 nm blue light. In the dark the pZF-HIS3 strain is a histidine auxotroph, while growth on media lacking histidine is conferred in blue-light only in the strain that also contains the ZDBD-CRY2/AD-CIB1 light-activated split transcription factor.

**Supplemental Figure 6:**
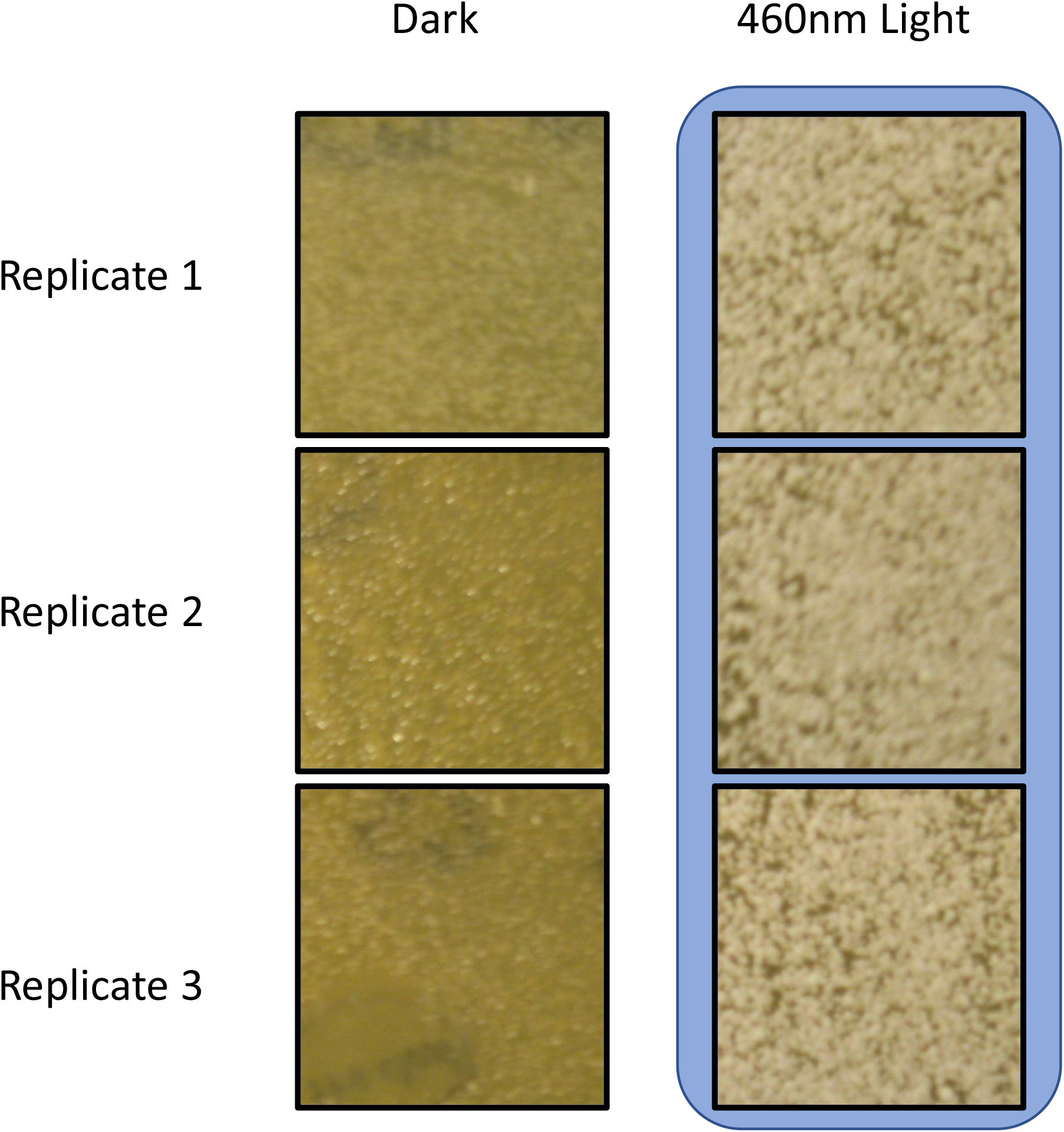
The pZF-SUC2 Yeast strain yMM1406 (Matα trp1Δ63 leu2Δ1 ura3-52 HO::SV40NLS-VP16-CIB1 loxP SV40NLS-Zif268DBD-CRY2PHR KanMX-pZF(3BS)-SUC2) was plated at a density of 0.119 and grown on YP-Sucrose in either 0 µW/cm^2^ or 57 µW/cm^2^ 460 nm blue light for 7 days (n=3 technical replicates per condition). Blue-light confers the ability to grow on YP-Sucrose, while the replicates grown in the dark show no discernable growth.

**Supplemental Figure 7:**
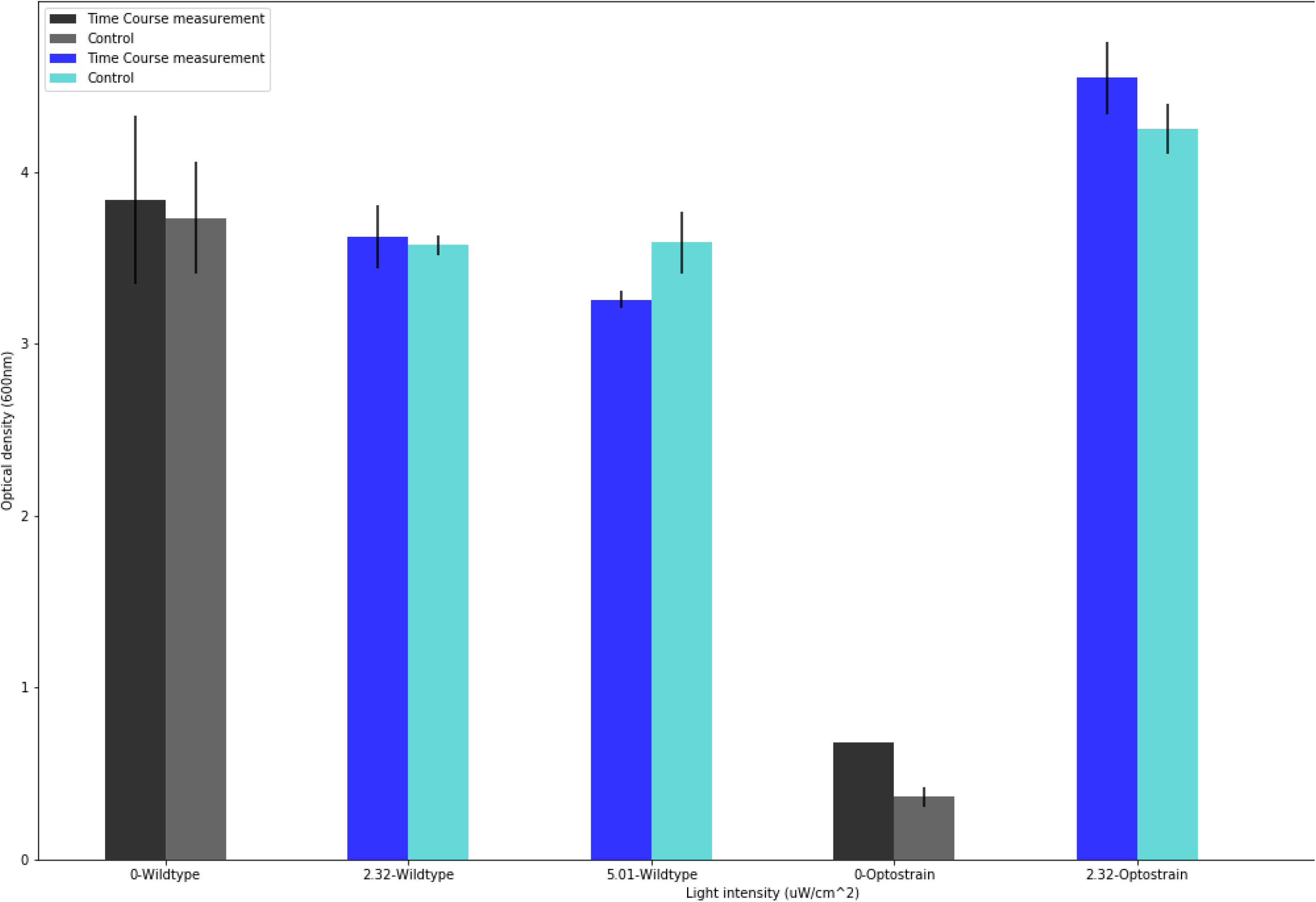
Endpoint point densities are compared between the time course sampled wells (black and blue bars) and control wells (gray and cyan bars) to determine whether there was any effects on the cultures upon the unintended addition of light during sampling for the time course experiment (**Figure 3**). The bars represent the mean density of the cultures at 52 hours for two technical replicates. The error bars represent the standard deviation from the population mean. Where there is no error bar (0, pZF-SUC2) the two measurements of density were the same value. A two-way ANOVA identified significant differences between the effect of strain and light intensity on density of the cultures (F, (3,6) = 46.72, p = 0). However, a multiple comparisons test did not identify any significant differences between the density reached for the control measurement and the time course measurement pairs for each intensity.

**Supplemental Figure 8:**
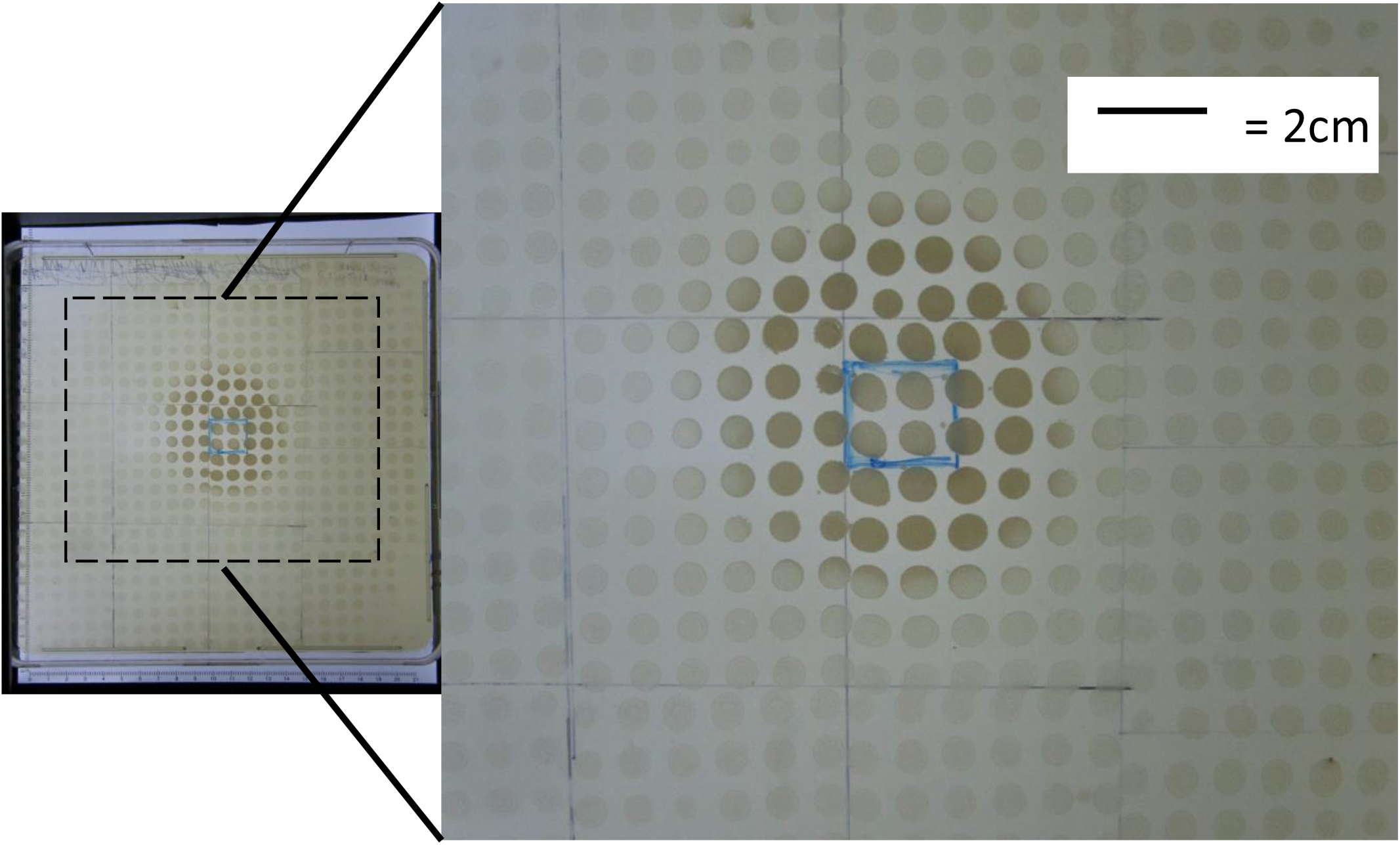
Spatial control of the optogenetic strain (yMM1406, Matα trp1Δ63 leu2Δ1 ura3-52 HO::SV40NLS-VP16-CIB1 loxP SV40NLS-Zif268DBD-CRY2PHR KanMX-pZF(3BS)-SUC2). The optogenetic strain was stamped on the surface of a 224 mm^2^ square petri dish, and only a small, 20 mm^2^ region (blue square) of the petri dish surface area was illuminated during incubation. The area around the illuminated portion of the plate (dashed outline) is magnified. A large number of colonies outside the illuminated region were able to grow.

**Supplemental Figure 9:**
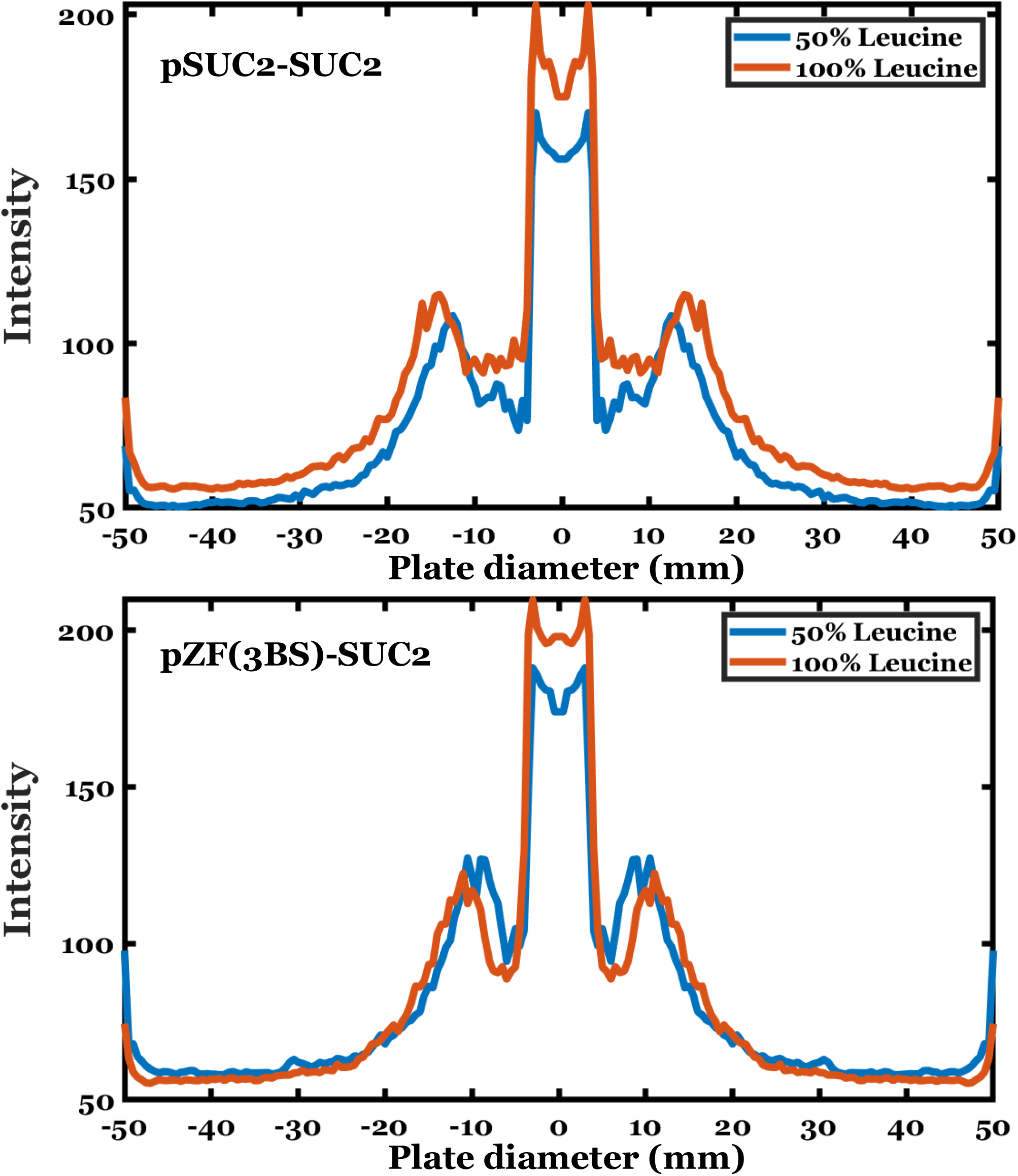
Plots represent an averaged radial intensity profile of the spotted plates across the diameter of the plate. The wild-type pSUC2-SUC2 (top) and pZF-SUC2 (bottom) are separately plotted and the two concentrations of leucine are distinguished by color (blue, 50% leucine; orange, 100% leucine). The wild-type pSUC2-SUC2 has a larger region of inhibited growth than pZF-SUC2 in both growth conditions.

**Supplemental Figure 10:**
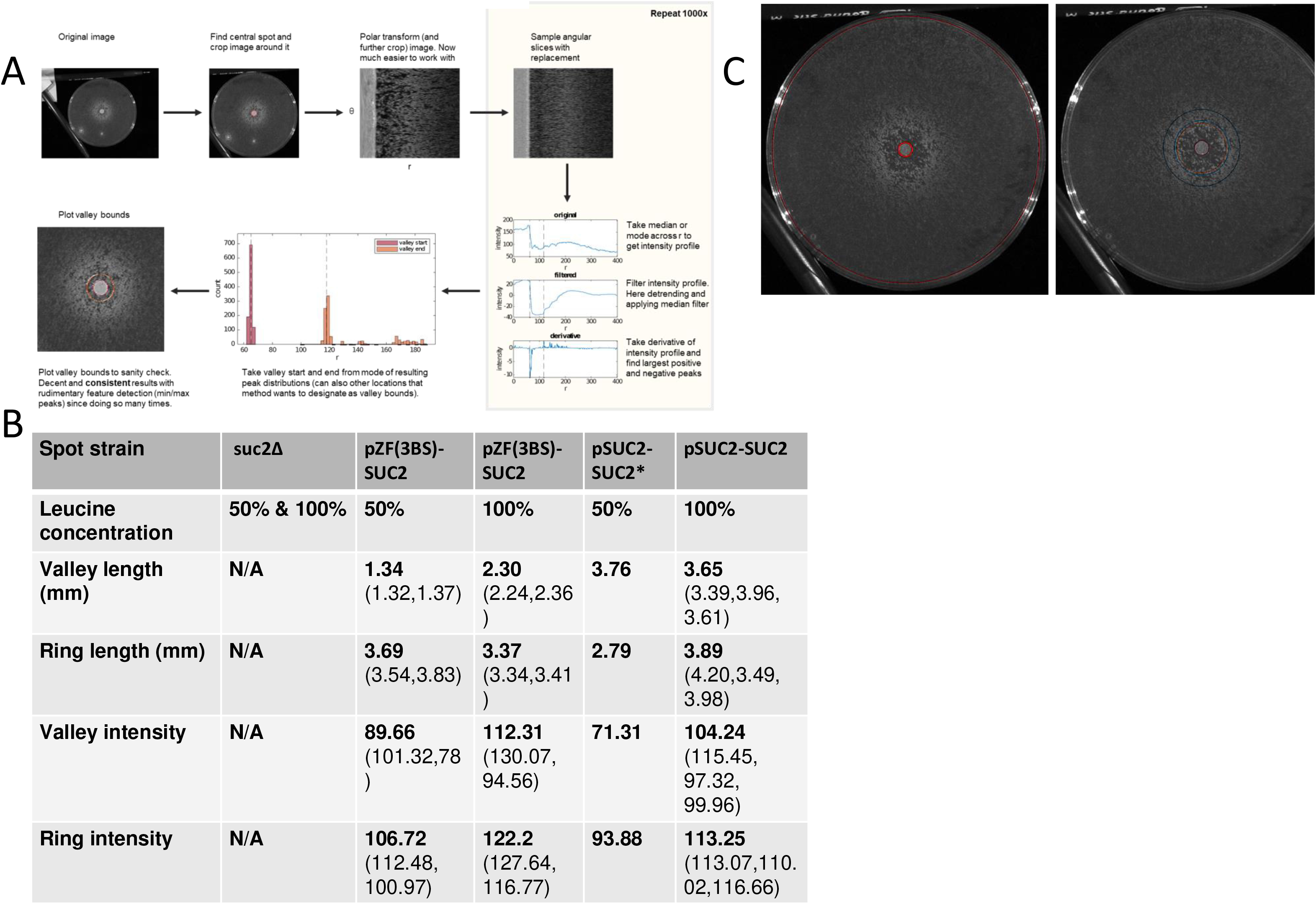
**(A)** Determination of pattern features and growth from plate images using a custom Matlab script. **(B)** Plate features of the emergent pattern between strains and between leucine concentration. Mean values are in bold, followed by both replicate measurements (*, n=1). **(C)** Representative image of a patterned plate. Inset highlights pattern regions, the spot (red), valley (orange) and ring (blue) quantified by the Matlab script.

